# Sequential PIDD1 auto-processing is essential for ploidy control in the liver and heart

**DOI:** 10.1101/2025.06.23.660994

**Authors:** Felix Eichin, Valentina C Sladky, Matthäus A. Reiner, Marina Leone, Ernesto Abila, André F. Rendeiro, Ralph Böttcher, Maik Dahlhoff, Thomas Kolbe, Andreas Villunger

## Abstract

Polyploidization refers to the balanced increase in gene copy number and is a feature of specialized cells in different mammalian tissues, including the liver and the heart. During organogenesis, hepatocytes and cardiomyocytes undergo scheduled polyploidization events to increase their cellular or nuclear DNA content. This is thought to improve cellular output and enable for rapid genetic adaptation in response to stress. Yet, excessive increases in ploidy can also be disadvantageous and increase the risk of genome instability. Hence, a dedicated machinery, the PIDDosome multi-protein complex, has evolved to prevent exacerbated increases in DNA content. Using targeted mutagenesis in mice, we show that the PIDDosome controls hepatocyte ploidy in a cell-autonomous manner and that sequential and quantitative auto-processing of PIDD1 is key for accurate control of ploidy in postnatal development of liver and heart. Stoichiometric imbalances in bioactive PIDD1-fragments impair p53-dependent and independent cell cycle arrest responses during organogenesis, as well as caspase-2-dependent apoptosis caused by centrosome amplification. Strikingly, targeted mutagenesis of the caspase cleavage motif in the critical E3-ligase controlling p53 protein levels, *Mdm2,* impairs ploidy control in hepatocytes, but not in cardiomyocytes, indicative of the existence of alternative caspase-2 substrates that help to restrict ploidy in the heart.

## Introduction

A fundamental task for a cell is to ensure that before division, it duplicates its DNA content precisely once and distributes it equally to daughter cells to maintain a stable, usually diploid, genome. Changes in chromosome numbers, such as the gains or losses of individual chromosomes or structural aberrations, are a hallmark of aneuploidy and can be observed in various diseases, most notably cancer^1^. In contrast to aneuploidy, polyploidy describes the balanced increase of the entire genome and whole genome duplication (WGD) is commonly seen in plants and some animal species but does not occur at the whole organism level in mammals^2–4^. However, some specific cell types in our body amplify their entire genome during differentiation and become polyploid, as seen in cardiomyocytes and hepatocytes^5–7^. The reasons for this are not completely understood but it is thought to help these tissues to cope with the metabolic and mechanical workload and allow swift adaptation to genotoxic stress^8^. Polyploidization can occur by several means; cells can undergo endoreplication and not even enter, or abrogate mitosis, or enter mitosis but skip cytokinesis^9^. Alternatively, cells can fuse, as seen in osteoclasts or macrophage giant cells, resulting in polyploid cells with multiple nuclei^10^. A critical secondary consequence of polyploidization is the accumulation of extra centrosomes^11,12^.

A key signalling pathway that limits the degree of polyploidization is induced by the PIDDosome, a multi-protein complex consisting of three different proteins, PIDD1, RIP-associated protein with a death domain (RAIDD/CRADD) and caspase-2. The P53-Induced Death Domain Protein 1 (PIDD1) is produced as a precursor protein that undergoes auto-processing at two conserved sites via an intein-like mechanism and thus generates three fragments, PIDD1-N, PIDD1-C and PIDD1-CC ^13,14^. Two of these, PIDD1-C and PIDD1-CC, can form different multi-protein complexes involved in distinct cellular signaling processes, namely the NEMO (NF-kappa-B essential modulator) PIDDosome and the caspase-2 PIDDosome, respectively^13–15^. Little is known about the role of PIDD1-N in downstream signaling. DNA damage was long thought to be the only stimulus promoting the formation of these complexes, resulting in cell death via caspase-2 activation or cell survival via nuclear factor kappa-light-chain-enhancer of activated B cells (NF-kB) signaling through the NEMO PIDDosome^13,15,16^. More recently, several groups, including ours, have established a link between PIDD1 and extra centrosomes as the activation cue for the formation of both signaling complexes^17–20^.

The caspase-2 protease is a highly conserved initiator caspase that becomes activated through dimerization within the PIDDosome^21^. Only a few proteins have been firmly established as substrates of active caspase-2, namely the E3 ligase MDM2, the BCL2 family member BID, and the ER resident site-1-protease, S1P^21–23^. The cleavage of BID into tBID induces apoptosis by promoting mitochondrial outer membrane permeabilization (MOMP). Proteolytic inactivation of MDM2, the master regulator of the tumor suppressor p53, results in p53 accumulation and, thus the transcription of p53 target genes, including cyclin-dependent kinase inhibitor 1 (*CDKN1A/p21*) ^22,24,25^. The PIDDosome/p53/p21 axis functions to induce a cell cycle arrest in cells that accumulate extra centrosomes, e.g., after cell division failure, thereby limiting the polyploidization of hepatocytes and cardiomyocytes in mice^18,26^. In addition, a molecular switch has been described by which FANCI, a member of the Fanconi anemia complementation group family, recruits PIDD1-CC in response to failed repair of interstrand DNA crosslinks to induce PIDDosome dependent cell death, which requires PIDD1 phosphorylation and SUMOylation^27,28^. Along similar lines, the induction of single strand DNA breaks during S-phase in response to treatment of cancer cells with topoisomerase 1 inhibitors reportedly promotes PIDDosome formation, placing caspase-2 at the intersection of cell cycle and cell death responses^29,30^.

In the NEMO-PIDDosome, PIDD1-C interacts with receptor-interacting serine/threonine-protein kinase 1 (RIPK1) and NEMO, facilitating the inhibition of the inhibitor of nuclear factor kappa B (IκBα) and thus activation of NF-κB^14,20^. Active NF-κB translocates to the nucleus and promotes cell survival and transcription of inflammatory genes and chemokines that facilitate recognition of cells with extra centrosomes by the innate immune system^20^. Of note, PIDD1-C has also been implicated in the regulation of DNA replication fidelity by interaction with PCNA, replacing p21 and thereby facilitating DNA synthesis across UV-induced DNA lesions^31^.

The regulation of the processing of full-length PIDD1 protein (PIDD1-FL) is not well understood. Early work by the Tschopp lab showed that PIDD1 interacts with several chaperones, including heat shock protein 90 (HSP90), which are essential for efficient auto-processing^32^. Furthermore, they found that substitution of serine 446 or 588 with alanine in the auto-processing motif (HFSW) of human PIDD1-FL impairs self-processing at these sites^14^. Further evidence indicates that PIDD1-FL can be directly processed into PIDD1-CC in cells, bypassing the PIDD1-C intermediate^14^. In addition, only PIDD1-CC but not PIDD1-C can efficiently form the caspase-2 PIDDosome together with the adaptor protein RAIDD, even though both fragments carry the C-terminal death domain (DD) that is critical for RAIDD binding^33^. Interestingly, sensing of extra centrosomes via the distal appendage protein ANKRD26, found at mother centrioles, requires PIDD1-FL protein, as none of its processed fragments can localize there^34,35^. Together, these observations suggest a highly complex mechanism regulating the availability and abundance of individual PIDD1 fragments required for signaling under various stress conditions, including incomplete cell division and different types of DNA damage.

In this study, we generated a panel of mouse strains harboring point mutations in the *Pidd1* gene to elucidate the importance of PIDD1 auto-processing for PIDDosome-dependent signaling events *in vivo* and validated the relevance of p53 as the key caspase-2 effector in ploidy control by targeted mutagenesis of D359 in the *Mdm2* locus.

## Results

### The PIDDosome controls hepatocyte polyploidization in a cell autonomous manner

Our previous study showed that the PIDDosome is important to limit the polyploidization of hepatocytes in mice. As all mouse mutants studied were full-body knockouts of individual PIDDosome components, it was unclear if the phenomena observed during liver development and regeneration were strictly cell autonomous^26^. Hence, we investigated liver ploidy in newly generated *Pidd1^f/f^* and *Raidd^f/f^* mice and compared them to *Casp2^f/f^*mice^36^ expressing *Cre* recombinase under the control of the *Albumin* promoter that restricts expression to hepatocytes (Fig. 1A). Flow cytometry-based ploidy analysis of hepatocytes isolated from adult animals revealed that *AlbCre* mice harboring floxed alleles of different PIDDosome components indeed showed increased liver ploidy, comparable to that observed in full-body *Pidd1* deficient mice. This demonstrates that ploidy control exerted by the PIDDosome in the liver is cell autonomous (Fig. 1B-D, Suppl. Fig. 1).

**Figure 1.**
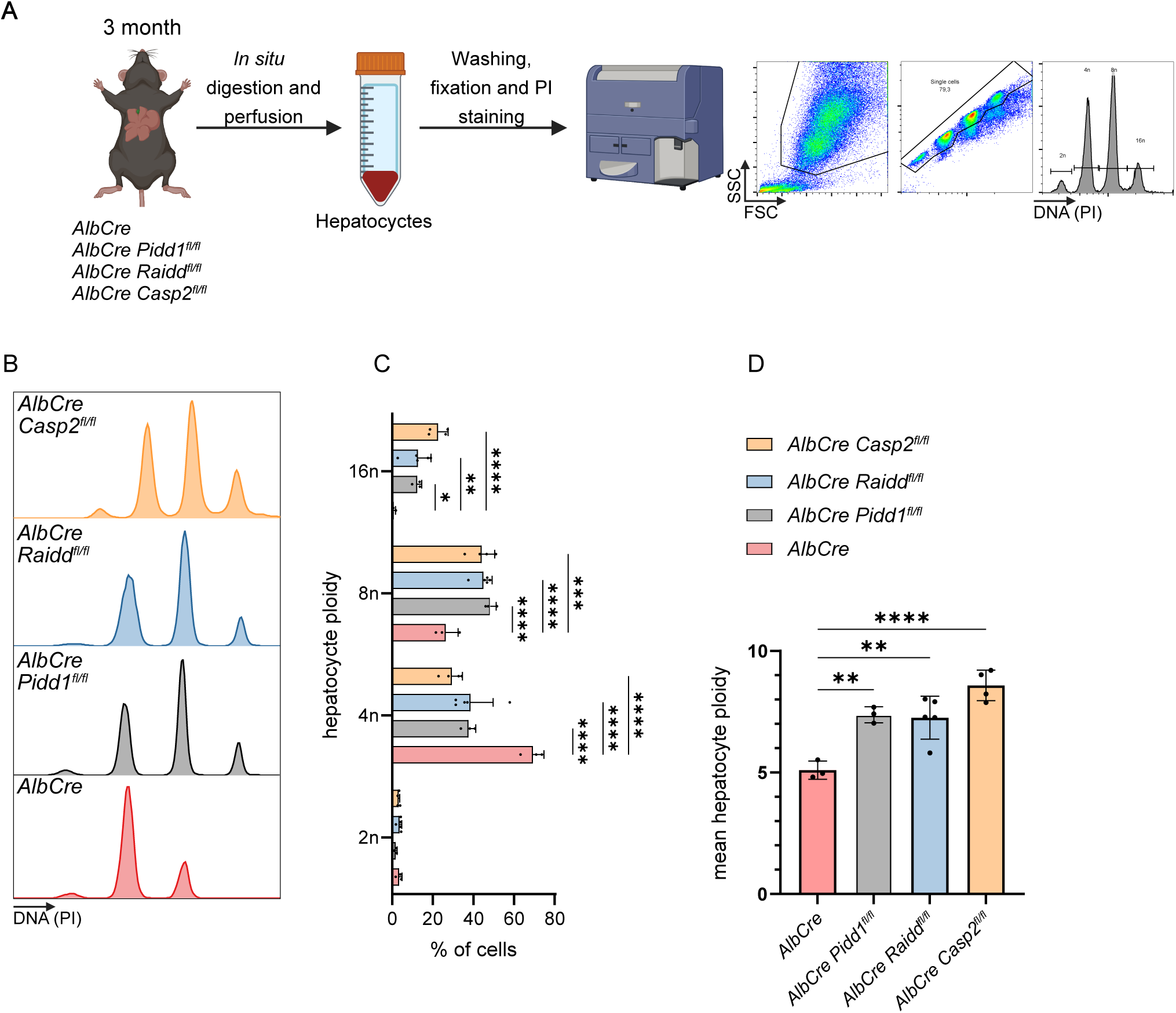
The PIDDosome limits hepatocyte polyploidization in a cell autonomous manner. A) Hepatocytes were isolated from 3-month-old mice by perfusion, followed by propidium iodide staining to measure their ploidy by flow cytometry. B) Representative histograms of the DNA content analysis of hepatocytes lacking individual PIDDosome components in the liver upon *AlbCre*-mediated deletion of conditional alleles; C) the quantification of the different hepatocyte ploidy fractions and D) their calculated mean ploidy. N=3-4. Statistical analysis: One-way ANOVA (for D) or two-way ANOVA (for C) Dunnett’s multiple comparisons. *=p<0.05, **=p<0.01, ***=p<0.001, ****=p<0.0001

### Precise PIDD1 auto-processing is critical for ploidy control during organogenesis

Among mammals, PIDD1 is very well conserved, especially around its two processing sites, indicating that Ser451 and Ser593 of the mouse protein are the counterparts of the reported auto-processing sites in human PIDD1 at positions Ser446 and Ser588, respectively (Fig. 2A).

**Figure 2.**
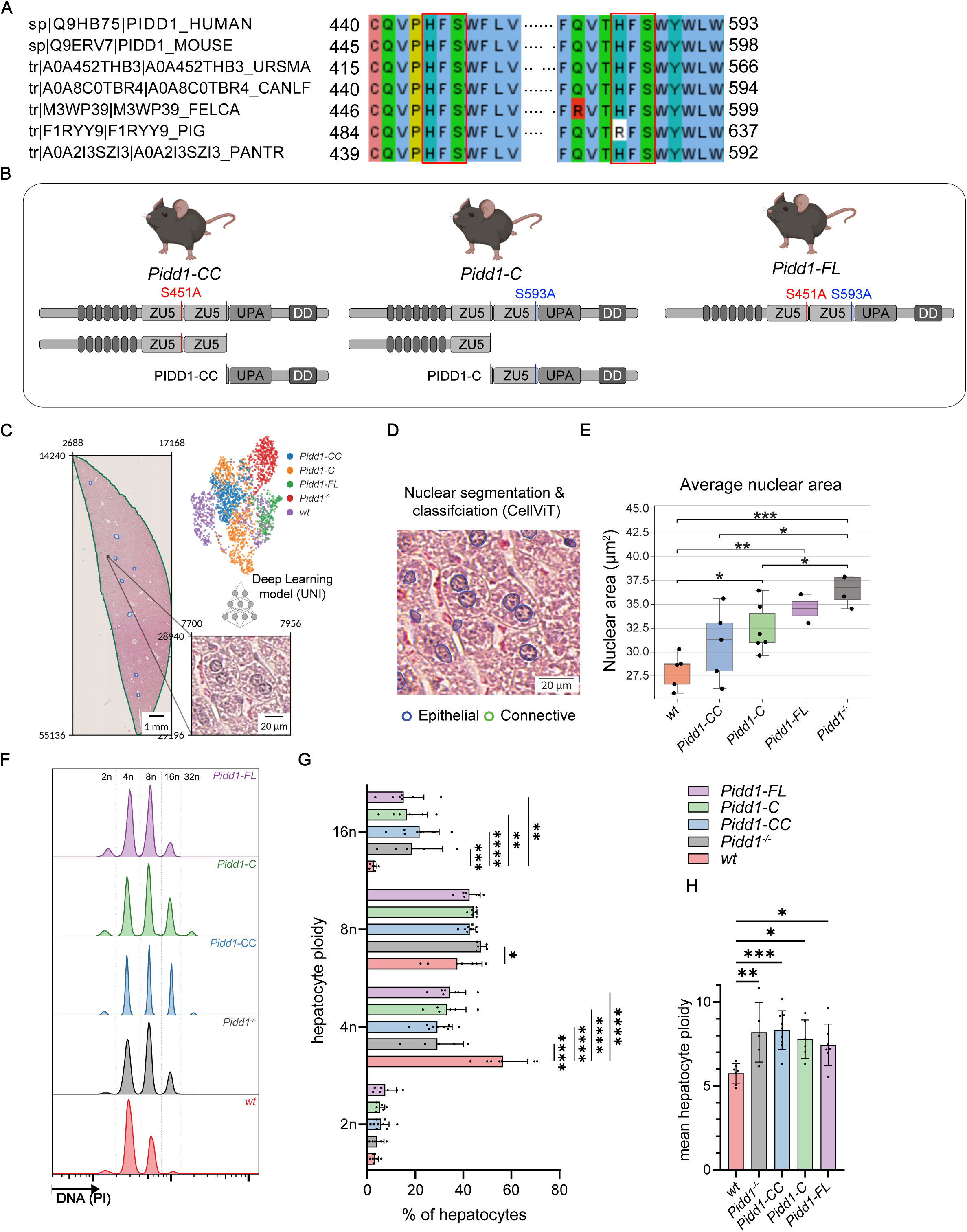
PIDD1 auto-processing mutants fail to limit hepatocyte polyploidization. A) Clustal omega alignment of a selection of mammalian PIDD1 amino acid sequences flanking both its auto-processing sites. B) Scheme indicating mutations in the *Pidd1* mutant mice and the PIDD1 proteins they are expected to produce. C) Representative H&E liver section of 3-months-old mice; schematic of tile-based morphological feature extraction, and UMAP embedding colored by genotype. D) Example tissue tile showing CellViT nuclear segmentation and classification. E) Quantification of mean nuclear area per slide (2-6 mice per genotype). F) Representative histograms of the DNA content analysis of hepatocytes isolated from 3-month-old mice evaluated by flow cytometry. G) Quantification of the different ploidy fractions H) Calculated mean ploidy of the hepatocytes. N=4-9. Statistical analysis: One-way ANOVA (H) or two-way ANOVA (G) Dunnett’s multiple comparisons. *=p<0.05, **=p<0.01, ***=p<0.001, ****=p<0.0001

To study the consequences of impaired PIDD1 auto-processing in organogenesis and create separation-of-function variants, we used CRISPR/Cas9 genome editing to mutate mouse embryonic stem cells (mESCs). These were used to generate mice carrying serine-to-alanine substitutions at position 451 (critical to create PIDD1-C), at position 593 (critical to create PIDD1-CC), or at both auto-processing sites, predicted to give rise to non-cleavable PIDD1-FL (Fig. 2B). For simplicity, we named gene-modified animals according to the protein they are expected to express. Mice expressing the S451A mutant version of *Pidd1* are expected to produce PIDD1-CC, while the S593A mutant mice are expected to express PIDD1-C, and the S451A, S593A double mutant only PIDD1-FL. Evaluation of litter size, body weight, sex distribution and allele frequency, no significant differences were observed for the PIDD1-C and –CC mice, compared to wildtype. Yet, we noted that females were underrepresented in the PIDD1-FL mutant strain, with fewer animals than expected expressing the mutated alleles in a homozygous state (Suppl. Fig. 1 A+B). Overall, we did not observe any gross anatomical or structural abnormalities in major organs. However, a deep learning-driven histological analysis of the liver sections showed morphological distinctions among the mouse genotypes (Fig. 2C).

Further, applying CellViT for nuclear segmentation and classification revealed that all three *Pidd1* mutant mouse strains possessed larger nuclei compared to wild-type animals, consistent with an increase in ploidy (Fig. 2 C-E, Suppl. Fig. 2C). Hence, we asked if impaired maturation of PIDD1 affects ploidy control in the liver in a quantitative manner. We anticipated that animals expressing PIDD1-FL or PIDD1-C would resemble *Pidd1^-/-^* mice, while PIDD1-CC expressing mice should show normal hepatocyte ploidy, comparable to wild-type animals. Thus, we isolated hepatocytes from 3-month-old mutant mice and the corresponding littermate controls and stained cells with propidium iodide to measure their DNA content by flow cytometry (Fig. 2F). Unexpectedly, all three mouse mutants showed a significant increase in ploidy, which strongly resembles the hyperpolyploidization noted in *Pidd1^-/-^* mice (Fig. 2F-H). The increase in ploidy seen in PIDD1-CC mice was unexpected, as these mouse mutants should be able to produce a functional PIDD1-CC fragment, allowing caspase-2 activation to limit hepatic polyploidization.

Next, we wanted to determine if this is a hepatocyte-specific phenomenon. Therefore, we also assessed the ploidy of cardiomyocytes (CM), another cell type known to limit polyploidy in a PIDDosome-dependent manner^38^. To do so, nuclei from snap-frozen hearts were isolated and their nuclear DNA-content determined using PCM1 as a marker that identifies CM nuclei, along with PI staining ^39^ (Fig. 3A). Again, PIDD1-FL and PIDD1-C mutant mice showed a strong increase in ploidy and with an about two-fold higher fraction of tetraploid nuclei compared to wild-type animals (Fig. 3B+C). In contrast to the liver phenotype, the PIDD1-CC mice showed an intermediate increase in ploidy, indicating a hypomorph phenotype (Fig. 3B+C, Suppl. Fig. 3A).

**Figure 3.**
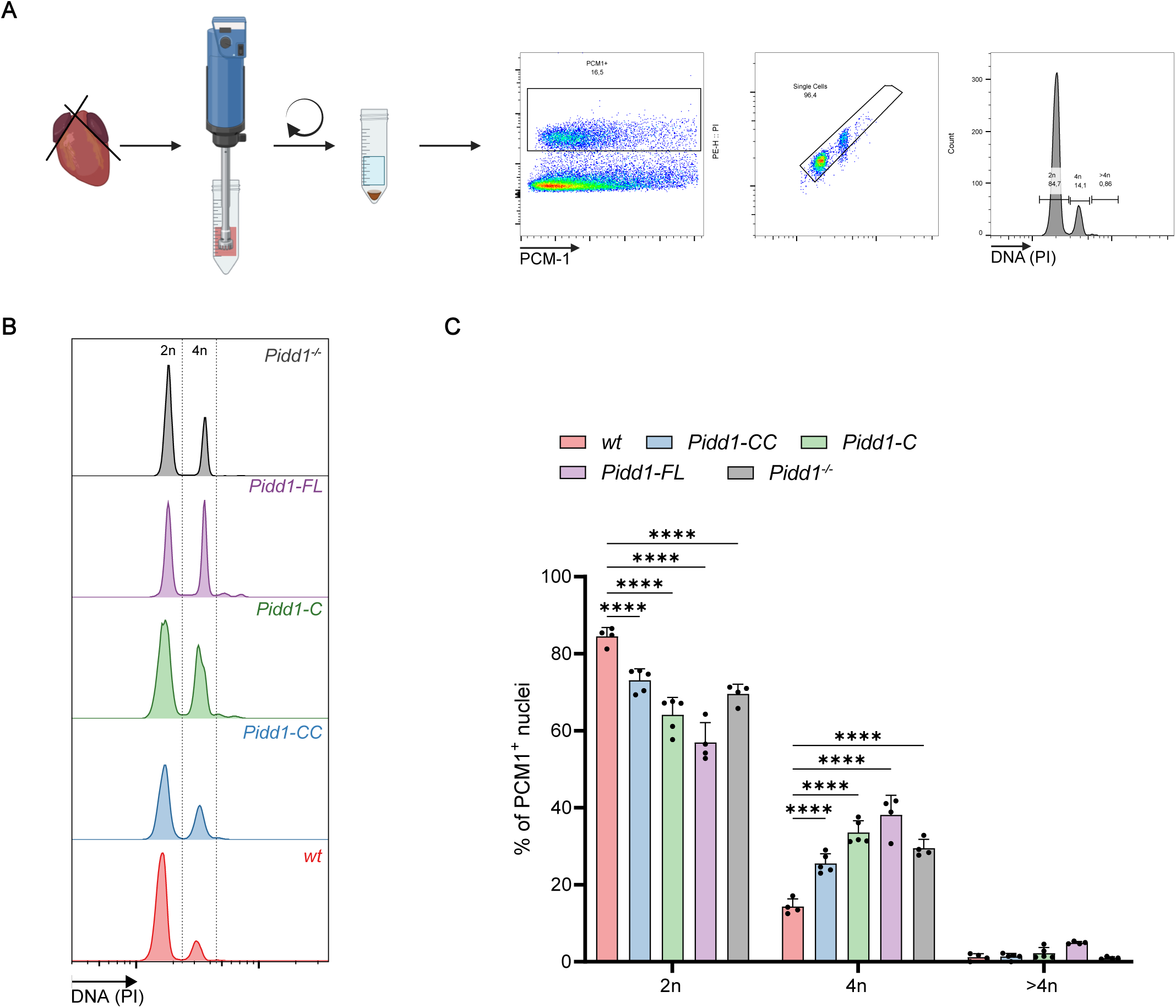
PIDD1 auto-processing mutants fail to limit the developmental ploidy increases of cardiomyocytes. A) Nuclei were isolated from frozen left and right ventricles of 3-month-old mice and cardiomyocytes (CM) ploidy was analyzed by flow cytometry using propidium iodide. PCM-1 co-staining was used to identify CM nuclei B)+C). N=4-5. Statistical analysis: Two-way ANOVA Dunnett’s multiple comparisons. *=p<0.05, **=p<0.01, ***=p<0.001, ****=p<0.0001

### Efficient production of PIDD1-CC depends on efficient auto-processing of Ser451

Since the PIDD1-CC mice showed a strong increase in hepatic and an intermediate increase in cardiac ploidy, we wondered whether (1) the mutation affects expression levels or (2) the direct processing of PIDD1-FL into PIDD1-CC is inefficient. To investigate the effects of PIDD1 processing in more detail, we isolated primary mouse embryonic fibroblasts (MEFs) from E14.5 embryos and generated lymphoid progenitor cells from young mice by transduction of bone marrow with an *HoxB8* encoding retrovirus^40^.

First, PIDD1 protein levels were assessed in MEFs by western blotting to determine if they could produce the expected fragments. We noted that MEFs derived from PIDD1-CC embryos failed to produce detectable amounts of the PIDD-CC fragment, yet exhibited increased levels of unprocessed full-length protein (Fig. 4A). In contrast, the PIDD1-C mutant cells generated the expected protein fragment that was not further processed to PIDD1-CC. Therefore, higher levels of PIDD1-C could be detected when compared to wild-type cells (Fig. 4A). As expected, the PIDD1-FL mutants could produce neither PIDD1-C nor PIDD1-CC. PIDD1-FL itself was expressed at very low levels.

**Figure 4.**
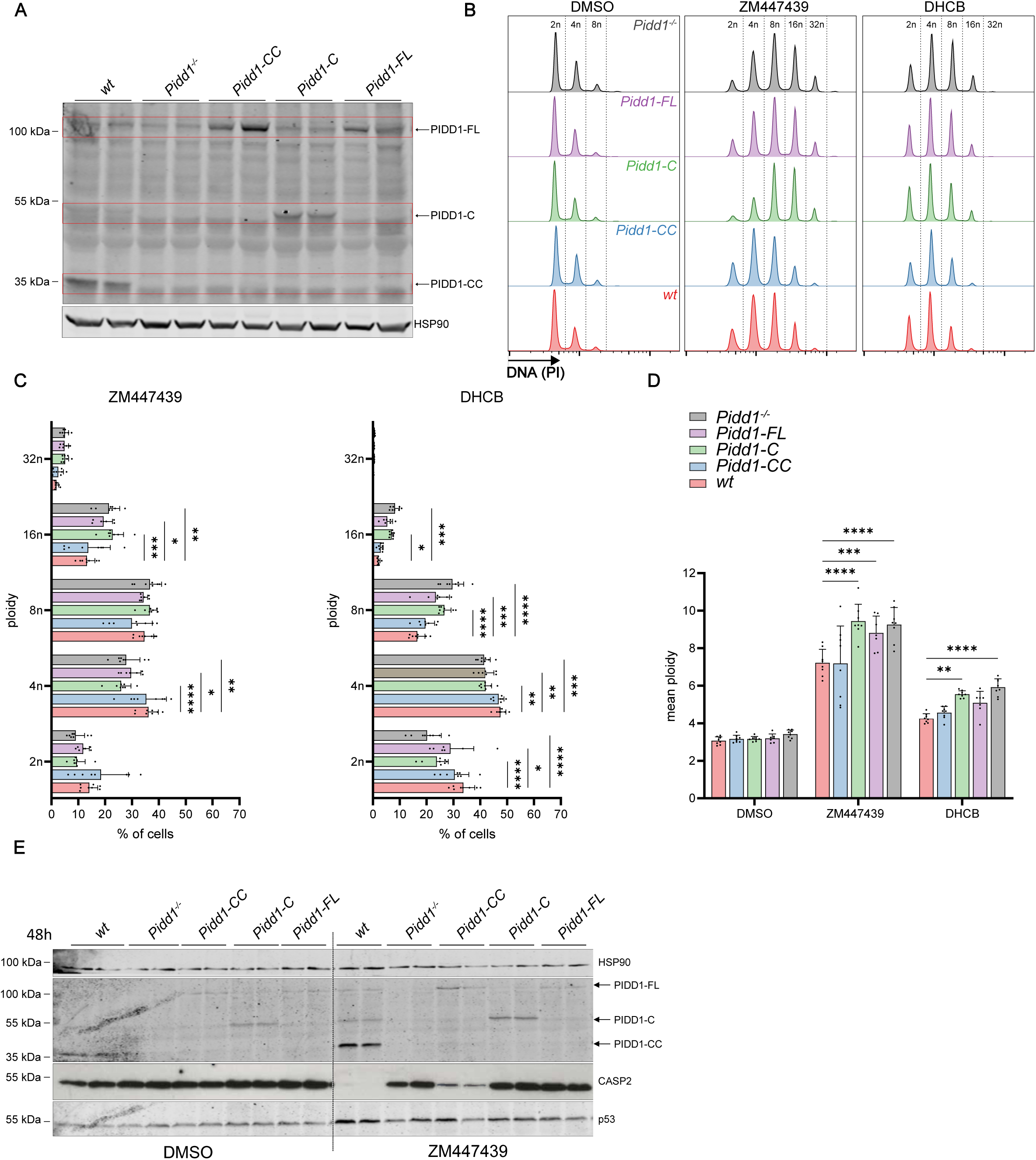
The S451A mutation impairs PIDD1-CC production but not PIDDosome activity in fibroblasts. A) Western blot of mouse embryonic fibroblasts (MEF) of the indicated genotypes. B-D) MEFs were treated for 48h with the Aurora kinases B inhibitor ZM447439 (ZM, 2µM) or dihydrocytochalasin B (DHCB, 2µM) to induce cytokinesis failure. Cellular ploidy was analyzed by propidium iodide staining and flow cytometric analysis. E) Biochemical analysis of PIDDosome activation in MEF. N=2 biological replicates and 3-4 technical replicates were generated. Statistical analysis: Two-way ANOVA Dunnett’s multiple comparisons. *=p<0.05, **=p<0.01, ***=p<0.001, ****=p<0.0001

As PIDDosome-mediated activation of caspase-2 is dependent on PIDD1-CC, very low expression levels could explain why the PIDD1-CC mice show increased ploidy in liver and heart. To test if these cells indeed fail to activate caspase-2 via the PIDDosome, early passages of primary MEFs were treated with an Aurora kinase B inhibitor (ZM447439) or dihydrocytochalasin B (DHCB) to induce polyploidization. These experiments indicate that primary MEFs do not arrest efficiently upon failed cell division, as wild-type cells become highly polyploid after 48 hours of drug treatment (Fig.4 B, C). Yet, clear differences could be observed between the genotypes: while the PIDD1-C and PIDD1-FL mutants behaved like PIDD1-deficient cells with an increase in ploidy, the PIDD1-CC mutant MEFs showed a similar ploidy increase as wild-type cells (Fig.4 B-D, Suppl. Fig. 4A-C). As the reduced ploidy of the PIDD1-CC MEFs suggested activation of the PIDDosome, we next analyzed hallmarks of pathway activation at the protein level. Western blot analysis showed a clear loss of pro-caspase-2 in ZM-treated wild-type and PIDD1-CC cells, suggesting pathway activation. In contrast, the other two mutants, like PIDD1-deficient cells, showed no loss of the caspase-2 pro-form at all. This clearly indicates residual activation of caspase-2 in PIDD1-CC mutant cells despite undetectable PIDD1-CC protein levels (Fig. 4E). In addition, a mild increase of p53 protein could be detected in these cells, although this was less obvious due to the high steady-state levels of p53 in primary MEF.

### The S446A mutation in PIDD1 strongly impairs auto-processing

To investigate further the impact of impaired auto-processing, we exogenously expressed the corresponding mutations in human PIDD1 in HEK 293T cells (Fig. 5A). Contrasting previous work^14^, the expression of human PIDD1 containing the S446A mutation resulted in a massive decrease in PIDD1-CC levels when compared to overexpressed wild-type PIDD1 (Fig. 5B). All other mutants behaved as expected, giving rise to PIDD1-C or PIDD1-FL only. Quantification of the different PIDD1 fragments detected using a PIDD-1-specific antibody revealed that ∼85% of the full-length protein remains unprocessed in the PIDD1-CC mutant with the S446A mutation. In contrast, in the wild-type and PIDD1-C overexpressing cells, unable to process further into PIDD1-CC, only ∼20% of unprocessed full-length PIDD1 protein was detectable (Fig. 5C). Similar results were obtained when conditionally re-expressing different PIDD1 variants in p53 proficient U2OS cells, depleted of endogenous *Pidd1* (Fig. 5D, E).

**Figure 5.**
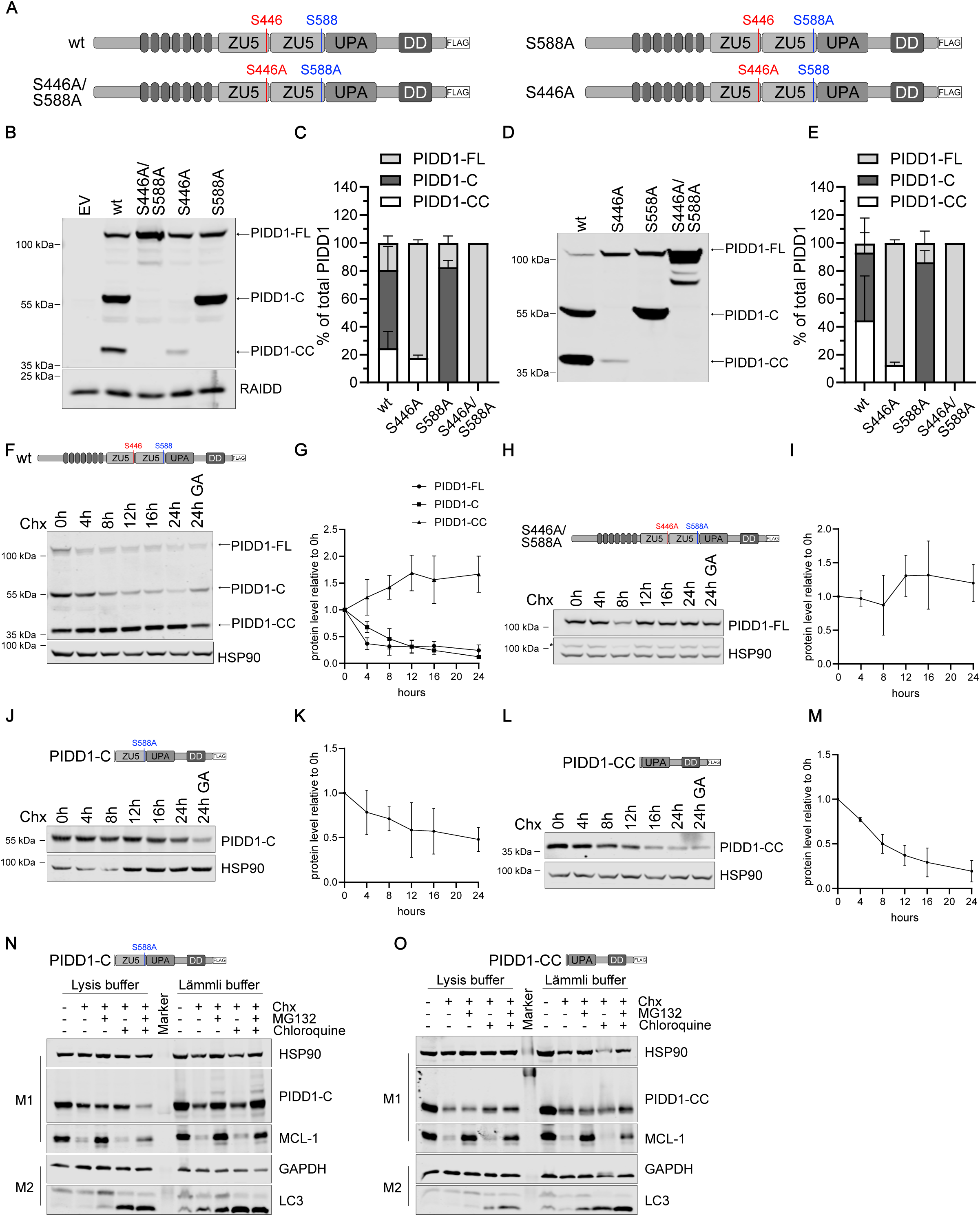
Analysis of PIDD1 auto-processing and fragment stability. A) Scheme of the different PIDD1 expression constructs used in B-D. B) Transiently transfected HEK 293T cells were analyzed two days after transfection for the presence of different PIDD1 fragments using the PIDD1-specific Anto-1 antibody. C) Quantification of B) (N=2, biological replicates). D) Flp-In T-REx™ U2OS cells were induced with Doxycycline for 2 days and E) expression of different PIDD1 variants was quantified by densitometric analysis, normalized to total PIDD1 signal (N=2, biological replicates). F-L) Puls-chase analysis of 293T HEK Flp-In™ T-REx™ cells lacking endogenous *Pidd1*, in which expression of the indicated PIDD1 variants was induced by the addition of Doxycycline for 36 hours. After washout, cells were treated with either DMSO, Cycloheximide (CHX), with or without the HSP90 inhibitor geldanamycin (GA) (N=3). N-O) Pulse-chase analysis of 293T HEK Flp-In™ T-REx™ cells lacking endogenous *Pidd1*, in which expression of the indicated PIDD1 protein variants was induced by the addition of Doxycycline for 36 hours. After washout, cells were treated with either DMSO, Cycloheximide (CHX), or treated with MG132 and/or Chloroquine. The cells were then lysed either in RIPA lysis buffer or boiled directly in Lämmli buffer.

### PIDD1-FL expression levels are solely defined by auto-processing efficiency

Based on these results, we could not determine whether reduced levels of PIDD1-CC in the S446A expressing human cells or our mouse mutant MEFs are solely due to impaired processing or to a reduced half-life of PIDD1-CC when PIDD1-C is not expressed. As these fragments were reported to co-IP upon PIDDosome activation in response to centrosome amplification^13^, they may influence their relative abundance due to stoichiometric needs, with excess proteins potentially being unstable and subject to degradation. To assess PIDD1 protein fragment stability, we performed a cycloheximide pulse-chase experiment with Flp-In-T-REx-293 cells lacking endogenous *Pidd1*. These cells allow doxycycline (Dox)-induced conditional re-expression of wild-type or mutant PIDD1 proteins. After induction of PIDD1 for 36 hours, followed by Dox-washout and addition of cycloheximide (CHX) to block translation, most of the full-length protein disappeared within the first four hours. In addition, PIDD1-C levels decreased rapidly, whereas PIDD1-CC appeared to be very stable (Fig. 5 F, G). However, since the fragments can be processed normally, the reduction in PIDD1-FL and PIDD1-C is likely to be due to auto-processing and not protein degradation, masking putative PIDD1-CC degradation. Additionally, addition of the HSP90 inhibitor geldanamycin, as published before^32^, reduced auto-processing to some degree.

Next, we expressed mutant PIDD1-FL, PIDD1-C or PIDD1-CC in Flp-In-T-REx-293 cells lacking endogenous *PIDD1* to further investigate their stability independent from auto-processing. This analysis revealed that PIDD1-FL protein unable to undergo self-processing is highly stable over time and is therefore most likely turned over under normal conditions solely by self-cleavage (Fig. 5H, I). Similarly, mutant PIDD1-C, unable to auto-process further, appears very stable and decreased by roughly 50% within 24 hours (Fig. 5J, K). In contrast, PIDD1-CC was degraded to a substantial extent over time (Fig. 5L, M). As previously reported, the interaction of HSP90 with PIDD1-C influences its stability. Thus, we inhibited HSP90 with geldanamycin, which resulted in decreased levels of PIDD1-C after 24 hours, as noted above (Fig. 5J). Next, investigated if the fragments are degraded via the proteasome or lysosome and therefore inhibited translation using cycloheximide (CHX) in the absence or presence with the proteasome inhibitor MG132, the lysosomal inhibitor chloroquine or both. Interestingly, PIDD1-C did clearly accumulate upon proteasomal inhibition, but this was only noted when cells were directly lysed in Lämmli buffer, but not under lysis conditions using detergents (Fig. 5N). In contrast, the degradation of PIDD1-CC was not slowed down by MG132 and/or chloroquine (Fig. 5O), suggesting aggregation in an SDS-insoluble fraction. Expression levels of the short-lived anti-apoptotic MCL1 and LC3B were monitored in parallel to confirm MG132 and chloroquine activity.

### PIDD1-CC expressing cells induce apoptosis upon cytokinesis failure

As we observed some differences with respect to ploidy in the liver vs. heart in PIDD1-CC mice, we also tested HoxB8-immortalized hematopoietic multipotent progenitor cells that were generated from the bone marrow of the mice for their response to forced polyploidization. In contrast to MEFs, these cells undergo apoptosis upon polyploidization, as caspase-2 preferentially cleaves and thereby activates the BH3-only protein BID in blood cells^25,41^ (Fig. 6A). Cytokinesis failure was induced using either ZM or DHCB. DNA content was assessed by propidium iodide staining and flow cytometric analysis, allowing quantification of ploidy and cell death using the fraction of cells with a DNA content smaller than that in cells in G1 phase (sub-G1) as a proxy for cell death. As seen in MEFs, the PIDD1-CC expressing hematopoietic progenitor cells behaved similarly to wild-type cells, with most cells dying within 24h upon ZM or DHCB treatment. In contrast, the PIDD1-C and PIDD1-FL expressing cells behaved like PIDD1-KO cells and survived cytokinesis failure significantly better (Fig. 6B, C, Suppl. Fig. 5A-C), confirming that PIDD1-CC is solely responsible for caspase-2 activation. Furthermore, we measured the ploidy increase in cells that were still alive at the time of analysis and detected a clear increase in PIDD1-FL and PIDD1-KO cells under both treatments (Fig. 6D, E), consistent with impaired caspase-2 activation upon centrosome amplification.

**Figure 6.**
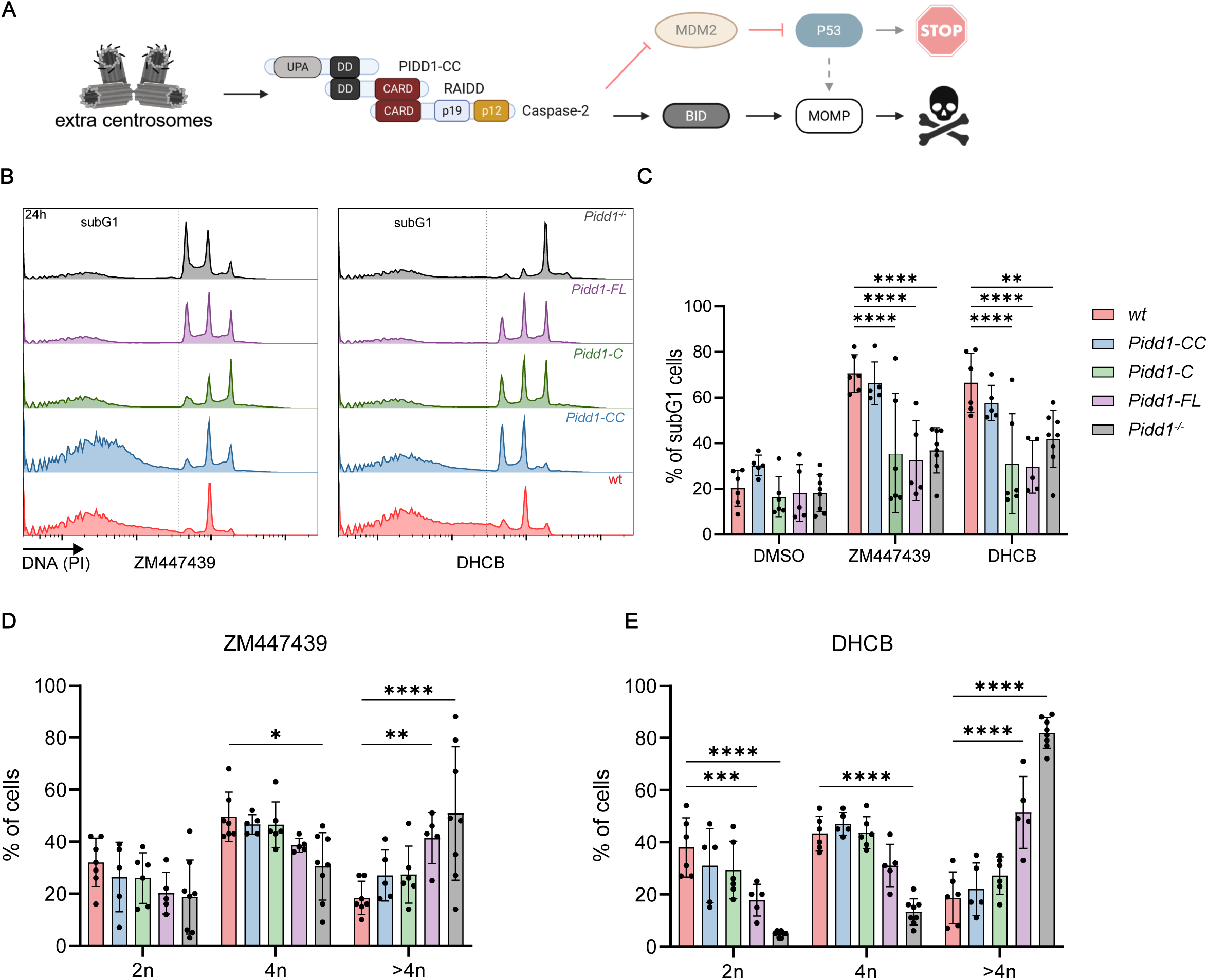
Cell death susceptibility of multipotent hematopoietic progenitor cells expressing different PIDD1 variants. A) Scheme of the established PIDDosome-driven and p53 dependent (shaded) or independent pathways of apoptosis induction in blood cells failing cytokinesis. B) Cell death and DNA-content analysis of HoxB8-immortalized hematopoietic progenitors with the indicated genotypes after exposure to either ZM447439 (ZM, 2µM) or dihydrocytochalasin B (DHCB, 5µM) for 24h. Representative histograms are shown. C) Quantification of B). N=2-3 biological replicates and 2-3 technical replicates. D+E) Ploidy analysis of B) after excluding the subG1 populations. Statistical analysis: Two-way ANOVA Tukey’s multiple comparisons. *=p<0.05, **=p<0.01, ***=p<0.001, ****=p<0.0001.

### Caspase-2 mediated cleavage of MDM2 is essential for hepatic ploidy control

The main pathway activated to induce a p21-dependent cell cycle arrest via the PIDDosome is the cleavage of MDM2 by activated caspase-2 separating the p53 binding domain from the RING domain of the E3 ligase (Fig. 7A)^22^. To test the universal action of this signaling axis in ploidy control, we also generated an *Mdm2* mutant mouse strain expressing an MDM2 protein resistant to caspase-2 cleavage. This was achieved by mutating aspartate D359, located in the established DVPD motif, recognized by caspase-2 (Suppl. Fig. 2A), to alanine, which is essential for caspase-2 mediated MDM2 cleavage^22^. We observed normal sex ratios and normal litter sizes in our cohort indicating that the mutation, that also distributes in a Mendelian ratio, does not affect general MDM2 function (Suppl. Fig. 6B), as a loss of *Mdm2* is embryonic lethal due to deregulated p53 expression levels *in utero* ^42,43^.

**Figure 7.**
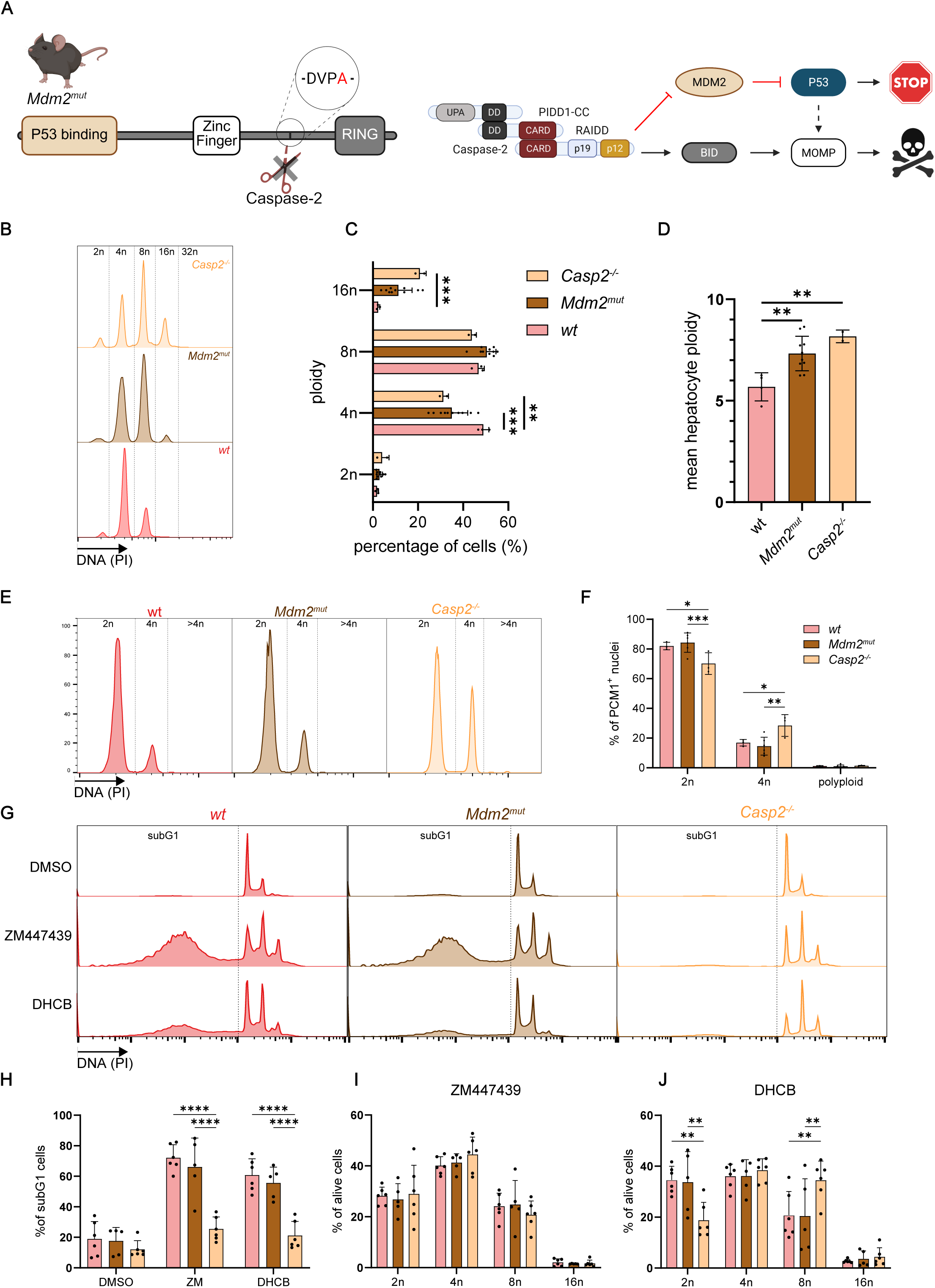
Caspase-2 mediated MDM2 processing is critical for ploidy control in the liver, but not the heart. A) Stick model illustration of MDM2 indicating the caspase-2 cleavage site, introduced into mouse ES cells by CRISPR/Cas9-based genome editing. B) Flow cytometric ploidy analysis of hepatocytes isolated from adult animals of the indicated genotypes. C+D) quantification of the ploidy fraction and the mean ploidy of the hepatocytes (N=2-10). E+F) Nuclei were isolated from frozen hearts of adult mice and their ploidy was analyzed by using the CM nuclei specific marker PCM-1 and flow cytometry (N=3-5). G) Cell death and DNA-content analysis of HoxB8-immortalized hematopoietic progenitors with the indicated genotypes that were treated with either ZM447439 (ZM, 2µM) or dihydrocytochalasin B (DHCB, 5µM) for 24h. Representative histograms. H Quantification of G) (N=3-4 biological replicates and 2 technical replicates). I+J) Ploidy analysis of I) after excluding the subG1 populations. Statistical analysis: One-way ANOVA (D) Two-way ANOVA (C, H-J) Tukey’s multiple comparisons. *=p<0.05, **=p<0.01, ***=p<0.001, ****=p<0.0001

As the *Mdm2* mutant animals should mimic PIDDosome deficiency, we first measured hepatocyte ploidy by flow cytometric analysis (Suppl. Fig. 6C). Hepatic polyploidy increased to a similar extent as observed in PIDD1 auto-processing mutants or caspase-2 deficient animals (Fig. 7B-D and Fig. 2B-D), confirming the proteolysis of MDM2 as the rate-limiting event in control of liver polyploidization. Next, we investigated the role of MDM2 in controlling cardiomyocyte polyploidization, as previous work from our laboratory indicated that this might not be dependent on p53^38^. Interestingly, the loss of caspase-2 mediated cleavage of MDM2 did not affect the nuclear ploidy of cardiomyocytes, as assessed in adult animals. Both wild-type controls and *Mdm2^mut^* mice had comparable levels of diploid and tetraploid CM nuclei. In contrast, ∼30% of the caspase-2 deficient CM were tetraploid and only 70% were diploid (Fig. 7E, F, Suppl. Fig. 6D), in line with our earlier findings^38^.

### MDM2 proteolysis by caspase-2 is not required for PIDDosome mediated cell death

Lastly, we generated HoxB8-immortalized multipotent hematopoietic progenitor cells from bone marrow of *Mdm2^mut^* mice to investigate the contribution of p53 activation via caspase-2 mediated cleavage of MDM2 to cell death after polyploidization. As before, we induced cytokinesis failure by treating the cells with ZM or DHCB. Flow cytometric ploidy analysis showed a similar fraction of wild-type and *Mdm2^mut^* blood cell progenitors dying in response to cytokinesis failure (Fig. 7G, H, Suppl. Fig. 6E, F). In addition, the surviving ZM-treated *Mdm2^mut^* cells showed no difference in ploidy compared to the controls (Fig. 7I, J). However, caspase-2-deficient cells were well protected from cell death under these conditions and became highly polyploid.

## Discussion

Polyploidization is a recurrent feature in mammalian tissues and cell types, yet of largely unclear significance^5^. The PIDDosome is known to limit the polyploidization of hepatocytes in mice during both postnatal liver development and in response to chronic or acute liver damage^26^. Importantly, this process appears to be conserved in the human liver and the lack of caspase-2 accelerates liver regeneration in a mouse model of partial liver hepatectomy^26^. This highlights the potential role of the PIDDosome and caspase-2 as a drug targets^26^. Therefore, understanding the mechanisms of PIDDosome auto-processing and the consequences of its impairment are vital. Although PIDD1 generates different fragments by self-cleavage, each with distinct functions, only *Pidd1* knockout mice have been studied so far, leaving consequences of impaired self-cleavage unexplored. To address this, we generated several *Pidd1* mutant strains that are deficient in auto-processing to study the function of the different PIDD1 fragments in a physiological setting. These mice developed normally and without any obvious abnormalities, consistent with the phenotype of *Pidd1^-/-^* mice^26^. However, our results demonstrate that direct generation of PIDD1-CC from the full-length precursor protein is insufficient to activate the PIDDosome during organogenesis *in vivo*, in contrast to earlier *in vitro* work^14^.

PIDD1-CC mutant mice that cannot process PIDD1-FL to PIDD1-C, fail to activate the PIDDosome in hepatocytes to limit polyploidization during organ development (Fig. 2F-H). However, we observed residual ploidy control in cardiomyocytes of PIDD1-CC mice and evidence of caspase-2 activation in MEFs following drug-induced cytokinesis failure. These organ and cell type specific differences may arise from different expression levels of PIDD1 that subsequently could result in higher PIDD1-CC levels in the heart compared to the liver. As even very low PIDD1-CC levels are sufficient for caspase-2 activation^17^, it is difficult to test such a hypothesis by western blotting, at least with the tools currently available. Alternatively, auto-processing may be affected by tissue or cell type dependent differences in the availability of chaperons, such as HSP90, reported to bind PIDD1 and affect autoprocessing^32^.

Our analysis of cell lines generated from the *Pidd1* mutant mice showed that the PIDD1-CC MEF could still activate caspase-2 to some degree, as indicated by a drop in its pro-form. This level of caspase-2 activation was still sufficient to limit ploidy increases in PIDD1-CC MEF after cytokinesis inhibition (Fig. 4D+E), suggesting that some cell types can respond very effectively even if upstream pathway activity is significantly compromised. Similarly, in immortalized hematopoietic progenitors, PIDD1-CC mutants induced robust cell death in response to forced polyploidization, whereas cells expressing PIDD1-FL or PIDD1-C did not, mimicking PIDD1-KO cells (Fig. 6B/C). This finding is consistent with the observation that cell death following cytokinesis failure is independent of the MDM2-p53 axis, but initially relies on caspase-2 processing pro-apoptotic BID into tBID^25^. Furthermore, PIDD1-CC mutant primary MEFs showed increased levels of PIDD1 full-length protein, while PIDD1-CC fragments remained undetectable by western blot (Fig. 4A). This suggests that direct processing is inefficient when Ser446 is mutated into alanine, yet trace amounts of PIDD1-CC are sufficient for PIDDosome activation. Of note, in some western blots, we could detect a very faint signal with the same molecular weight as the PIDD1-CC fragment seen in wild-type cells, indicative of residual auto-processing.

Our findings differ from those reported by Tinel et al. in 2007, who found that by mutating the first auto-processing site, a substantial amount of PIDD1-CC is still produced directly from the full-length protein in a highly efficient manner ^14^. Currently, we cannot explain the altered processing behavior observed *in vivo*, but our *in vitro* experiments using MEFs provide insight into the phenotype observed in mice. Although PIDD1-CC expressing cells retain partial caspase-2 activation, these mice fail to activate the PIDDosome efficiently *in vivo*. This discrepancy between *in vivo* and *in vitro* likely arises from the artificially high expression levels of exogenous PIDD1 protein in cell lines. We conclude that the strongly reduced processing ability of the S451A PIDD1 mutant in combination with its low expression *in vivo* results in insufficient amounts of PIDD1-CC for PIDDosome activation, likely affecting complex stoichiometry or dynamics of assembly.

In addition, we show that the turn-over of full-length PIDD1 appears to be regulated solely via auto-processing to PIDD1-CC, as mutant non-cleavable PIDD1-FL is extremely stable. Interestingly, the degradation of PIDD1-CC was not affected by inhibiting the proteasome or the autophagolysosomal pathway (Fig. 5 O), asking for follow-up studies into how PIDD1-CC is eventually turned over in cells.

Of note, in contrast to the PIDD1-CC mutant, the other two PIDD1 mutants behaved as expected: PIDD1-FL is not processed at all and PIDD1-C is not processed further into PIDD1-CC. Thus, these mutant mice are interesting tools for studying the function of PIDD1-FL or PIDD1-C *in vivo*. Although these animals showed no overt phenotypes in steady-state, challenging these mouse mutants, e.g., by inducing extra centrosomes using a *Plk4* transgene or by interfering with cytokinesis knocking down *Anillin,* could provide insights into whether the NEMO-PIDDosome ^20^ has an impact on liver regeneration or liver diseases by affecting sterile inflammation. These animals can also help to clarify if impairing PIDD1 auto-processing would provide an alternative strategy to inhibiting caspase-2. In addition, the proposed role of PIDD1-C in translesion DNA synthesis upon UV-induced DNA damage can now be investigated using the mice generated in this study^31^. Furthermore, it would be very interesting to investigate if the recently described SUMOylation-dependent activation of PIDD1 upon DNA damage is also required for PIDDosome activation by extra centrosomes *in vivo*^28^.

Finally, the increase in hepatic but not cardiac ploidy seen in *Mdm2* mutant mice strongly suggests that the PIDDosome restricts polyploidy in liver and heart by distinct caspase-2 substrates, rather than solely via the well-described MDM2/p53/p21 axis^18,26^ (Fig. 7C,F). Additional studies are needed to define new substrates in caspase-2 regulated ploidy control, or if activation of BID may suffice to clear cells after polyploidization in the heart.

## Material and Methods

### Generation of new mouse strains

#### Generation of new mouse strains

*Pidd1* floxed mice (B6;129-Pidd1^tm1Ozg^) were generated by *OzGene* flanking exon 4-10 with *LoxP* elements in C57BL/6N ES cells. Mice carrying a conditional *Raidd* allele (synomym: *Cradd*; C57BL/6-Cradd^tm1a(EUCOMM)Hmgu^) were created by injection of BALB/C blastocysts with gene targeted ES cells where *LoxP* elements were introduced in the *Cradd* locus upstream/downstream of exon 2 from the International Mouse Phenotyping Consortium. *Casp2^f/f^*mice (B6.Cg-Casp2^tm1.1Ejac^; Jax Stock #038519) were previously described^44^ and kindly provided by Etienne Jacotot, Universite Paris, Cite, Paris, FR.

The *Pidd1* mutant strains were generated by mutating KH2 mouse embryonic stem cells using CRISPR/Cas9 and injecting these cells into C57BL/6N blastocysts. In short, the KH2 cells were co-electroporated (Bio-Rad GenePulse Xcell Electroporator) with the pSpCas9(BB)-2A-GFP (PX458) vector, a gift from Feng Zhang (Addgene plasmid #48138)^45^, targeting the required locus (sgRNA_Pidd1_S451A: 5’CGACCAGGAACCAGGAGAAG3’, sgRNA_Pidd1_S593A: 5’ GGTCACGCACTTCTCCTGGT3’, sgRNA_Mdm2_D359A: 5’TCTGTCAGCTTTTTGCCATC3’) and a complementary ssDNA oligo repair template (ssDNA_PIDD1_S451A: 5’ TCCCCCGTGTTCCAGAGGCTCTGGGCTCGCTGCCAGGTcCCtCAtTTtgCgTGGTTtCTGGTCGTTTTACGCCCAGTATCCAACACTTGCCTTTTG3’, ssDNA_PIDD1_S593A: 5’GCACTGGAATTCACCCACCTGTATGCACGCTTCCAGGTtACcCAtTTtgCCTGGTCAGTGTCCCCTAGCTTCCGCAGGACCCCTTCTGCC3’, ssDNA_MDM2_D359A_ 5’ CAAACTGGAAAACTCAGCTCAGGCAGAAGAAGGCTTGGAcGTcCCaGccGGaAAAAAGCTGACAGAGAATGATGCTAAAGAGCCATGTGC3’, bases in lowercase letters indicate mutations). The repair templates contained the desired mutations and additional silent mutations for easier genotyping. The mutation sites were identified by protein sequence alignment of the human and mouse gene/protein sequences (Uniprot: PIDD1: Q9HB75 and Q9ERV7, MDM2: Q00987 and P23804; *Pidd1* Gene: NC_000073, Gene ID 57913, *Mdm2* Gene: NC_000076, Gene ID 17246) using the alignment tools of UniProt and SnapGene. GFP positive cells were sorted with a cell sorter (BD FACSAria™ III) 24h after electroporation and used to generate single cell clones that were first screened by PCR and then validated by Sanger sequencing. Validated single-cell clones were injected into C57BL/6N blastocysts and implanted into surrogate CD1 animals. Germline transmission was confirmed by genotyping PCR (S451A: frw: 5’CAGGTcCCtCAtTTtgCgT3’, rev: 5’CTGTGAAAGGCACAGCAGA3’, S593A: frw: 5’ GCATCAGTAAGCCCTCTGCT3’, rev: 5’ GGGACACTGACCAgGca3’, Mdm2_D359A: frw: 5’ ATGCTGAAAATTAAGCTACATGGT3’, rev: 5’CTCTGTCAGCTTTTTtCCgg3’; bases in lower cases indicate mutant specific bases) and Sanger sequencing. *Pidd1* mutant mouse strains were maintained on a mixed C57BL/6NxSV129 background.

### Ploidy measurement of hepatocytes

Hepatocytes from 3-month-old mice, indiscriminate of sex, were isolated as previously described ^26,46^. In short, the livers were immediately perfused *in situ* with 70 ml perfusion buffer (7 ml/min, 0.14MNaCl, 6.7mMKCl, 10mMHepes, 0.1mMEGTA, pH=7.4) after sacrificing the mice by cannulating the inferior vena cava and then with 40 ml collagenase buffer (66.7 mM NaCl, 6.7 mM KCl, 4.7 mM CaCl2*2H2O, 10 mM Hepes, pH=7.45, containing 25 mg/ml Liberase TM Roche, 2.5ml/min). Afterward, the livers were gently disrupted in cold DMEM medium (Sigma-Aldrich, D5671), and the hepatocytes were centrifuged 3 times for 3 min and 30 x g to clear away cell debris and other cells. The purified hepatocytes were fixed in ice-cold 70% ethanol and stored at −20°C before measuring their DNA content by propidium iodide staining (33.3 µg/ml PI, 0.01 µg/ml RNaseA in PBS) and a LSRII cytometer (BD Biosciences) or an Attune NxT flow cytometer(Thermo Fisher). As a small fraction of hepatocytes are diploid, we used the lowest DNA intensity peak to define the ploidy fractions. For the representative histograms, we aligned the peaks of the different samples for better visualization.

### Ploidy measurement of cardiac nuclei

Frozen hearts from 3-month-old animals, regardless of sex, without the atria, were cut into small pieces and then homogenized by a TP 18/10 Ultra-Turrax probe homogenizer (IKA, Germany) at 20 000 rpm for 20 seconds. Afterward, the cell suspension was passed up and down 8-10 times through a 20-gauge needle to break open the cells (the suspension was checked under the microscope for intact cells after 8 times). Then, the solution was passed through a 100 µm and a 70 µm strainer to get rid of intact cells and debris. After a centrifugation step of 10 min at 700 xg, the pellet was resuspended in a sucrose buffer and centrifuged for 60 min at 13,000 xg. The nuclei were then stained in nuclei storage buffer with rabbit anti-PCM1 antibody (1:300, HPA023370, lot. number: 000007967, Sigma-Aldrich) overnight, washed and stained for 1h with goat anti-rabbit Alexa 647-conjugated antibodies (1:500, Life Technologies). After washing with PBS, the nuclei were then incubated with propidium iodide and analyzed with a LSRII cytometer (BD Biosciences). For the representative histograms, we aligned the histograms so that the peaks of the different samples are easier to compare.

### Generation of mouse embryonic fibroblasts

MEFs were generated from 14.5-day-old embryos. In short, pregnant female mice of the indicated genotypes were sacrificed and the embryos were extracted. After dissecting the embryos, removing fetal liver and brain, the remaining tissue was minced into small pieces and incubated with trypsin for 20min at 37°C on a thermoshaker. Afterwards, the cell suspension was seeded onto a 15 cm cell culture dish and cultured in normal DMEM medium (Sigma-Aldrich D5671, 7 % FCS (Sigma-Aldrich F0804), 2 mM L-glutamine (Sigma-Aldrich G7513), 100 μg/ml streptomycin and 100 units/ml penicillin (Sigma-Aldrich P0781)) at 37°C and 5% CO2.

### Generation of HoxB8 cell lines from bone marrow

The multipotent hematopoietic progenitor cells were generated as previously described^40^. In short, the bone marrow of one femur of mice of the indicated genotypes was extracted and cultured for one day in Opti-MEM™ (Gibco, Cat. 31985047) supplemented with 10% FCS (Sigma-Aldrich F0804), 2 mM l-glutamine (Sigma-Aldrich G7513), 100 U/ml penicillin, 100 μg/ml streptomycin (Sigma-Aldrich P0781), 50 μM 2-mercaptoethanol, 10 ng/ml IL-3 (Lot #012248 K0623, PeproTech), 20 ng/ml IL-6 (Lot #112050 H0223, PeproTech). Then the cells were infected with the HoxB8 virus by spin infection (3x 30 min 380 xg at 37 °C, rigorously vortexing in between) and cultured afterwards with RPMI-1640 medium (Sigma-Aldrich R0883-500ML, 7 % FBS, 2 mM L-glutamine (Sigma-Aldrich G7513) and 100 U/ml penicillin) containing 1 µM ß-estradiol and 20% supernatant of FLT-3L expressing B16-melanoma cells.

### Cytokinesis failure experiments

MEFs from early passages 2-4 were cultured in normal DMEM medium with either DMSO, ZM447439 (2 µM, Selleckchem #S1103) or Dihydrocytochalasin B (2 µM, Sigma-Aldrich #D1641) for 48 hours and then harvested for protein extraction with RIPA lysis buffer and fixation with 70% ethanol for ploidy analysis as described for the hepatocytes with propidium iodide. The multipotent progenitor cells (HoxB8 cells) were treated with DMSO, ZM447439 (2 µM) or Dihydrocytochalasin B (5 µM) in RPMI-1640 medium containing ß-estradiol and FLT3 ligand for 24 hours and then fixed in 70% ethanol before ploidy analysis.

### Immunoblotting

Protein extraction was performed with a standard RIPA lysis buffer and protein concentration was measured with Bradford reagent to enable equal protein loading for western blot analysis. The membranes, both nitrocellulose and PVDF membranes, were blocked in 5% milk in PBS-T for 1h, followed by overnight incubation of the indicated primary antibodies (see table for clones and dilutions Table1). For protein detection, either a secondary antibody directly linked to HRP or a fluorophore (Table 1) was used and, after washing, membranes were exposed to a photo film using WesternBright™ ECL spray (Advansta #K-12049-D50), or the LICORbio Odyssey DLx Imaging System.

### Generation of stable and inducible cell lines

The Flp-In-T-REx-293 cells (CVCL_U427) were generated and described before^20^. Flp-In-T-Rex U2OS cells expressing the different PIDD1 protein variants, were generated by first deleting the endogenous *Pidd1* gene by CRISPR/Cas9 (LentiCRISPRv2, Zhang lab Addgene Plasmid #52961, sgRNA 5’GCCGATAGCGGATGGTGATG3’) and then co-transfection of the cells with a Flp-Recombinase plasmid (pOG44, Thermo Fisher) and the cDNA of the sgRNA-resistant PIDD1 variants cloned into pcDNA5 (Thermo Fisher, #V103320). Cells were selected for successful integration of the pcDNA5 vector using 150 µg/ml hygromycin B (Roth, #CP12.1) for one week.

### Protein stability assays

The expression of the PIDD1 constructs in the Flp-In-T-REx-293 cells were induced for 36 hours by the use of normal DMEM medium with 50 ng/ml doxycycline (Sigma-Aldrich, D9891). Afterwards, the medium was aspirated and cultured in medium without doxycycline containing cycloheximide (10 µM; Sigma-Aldrich #C1988) and/or, 1 µ/ml geldanamycin (Roth #HN71.1), 100 µM Chloroquine and 10 µg/ml MG132 (Sigma-Aldrich #C2211). The western blots where quantified using Image Studio Ver 5.2 (LICORbio) and normalized to the 0h time point to assess stability over time.

### PIDD1 processing assays

To analyze the processing efficiency of the different PIDD1 variants, we cloned different versions of PIDD1 cDNA into the pcDNA5 vector and then transfected into 293T HEK cells. Two days after the transfection, we harvested the cells and used protein lysates for western blot analysis. To assess processing efficiency in the Flp-In U2OS cells lacking endogenous PIDD1, cells were induced with 100 ng/ml doxycycline (Sigma-Aldrich, D9891) for 48h before harvesting. The western blot signal of PIDD1-FL, PIDD1-C and PIDD1-CC were quantified using Image Studio Ver 5.2 (LICORbio) and the relative share of all three fragments in relation to the total signal achieved was calculated.

### Deep learning-driven analysis of H&E liver sections

We processed mouse liver whole-slide histopathological images using CellViT for end-to-end nuclear segmentation and classification, yielding nuclear polygons and cell-type labels (e.g., epithelial)^37^. In parallel, we employed the LazySlide framework to segment tissue regions and tile each slide, then extracted high-dimensional image features from each tile using the deep learning vision model UNI^47,48^. From the CellViT-derived nuclear polygons, we calculated per-slide average nuclear area (µm²). We compared genotype-specific differences in nuclear area, using independent-samples t-tests with Benjamini–Hochberg FDR correction, annotating significant comparisons (p < 0.05) via the Statannotations Annotator^49^.

## Acknowledgements

We would like to thank Claudia Soratroi, Irene Gaggl and Julia Heppke for technical support, Sebastian Herzog and Yasmin Wolf for help with ES cell work, Martin Saurwein and Maria Fischer for animal care. We thank Prof. Reinhard Fässler for enabling ES cell work. We thank Mariana E G Araujo for providing us with the LC3B antibody. AV acknowledges support by the FWF (P36658, I6642), as well as the ERC, AdG 787171 (POLICE). FE acknowledges support from the DOC fellowship program of the Austrian Academy of Sciences (ÖAW). ML acknowledges support by the Austrian Science Fund (FWF) (Lisa Meitner, M 3115-B). For open access purposes, the author has applied a CC BY public copyright license to any author-accepted manuscript version arising from this submission.

## Author contributions

Conceptualization, AV, VCS, FE; Methodology, RB, TK, MD; Investigation, FE, MAR, ML, EA; Visualization, EA, FE; Supervision, AFR, VCS, AV; Writing— original draft, FE, AV; Writing—review and editing, VCS, TK, AFR, RB; Funding acquisition FE, ML, AV.

## References

1. Ben-David, U. & Amon, A. Context is everything: aneuploidy in cancer. Nat. Rev. Genet. 21, 44–62 (2020).

2. Van De Peer, Y., Mizrachi, E. & Marchal, K. The evolutionary significance of polyploidy. Nat. Rev. Genet. 18, 411–424 (2017).

3. Otto, S. P. The Evolutionary Consequences of Polyploidy. Cell 131, 452–462 (2007).

4. Otto, S. P. & Whitton, J. Polyploid incidence and evolution. Annu. Rev. Genet. 34, 401– 437 (2000).

5. Pandit, S. K., Westendorp, B., Bruin, A. De & De Bruin, A. Physiological significance of polyploidization in mammalian cells. Trends Cell Biol. 23, 556–566 (2013).

6. Gan, P., Patterson, M. & Sucov, H. M. Cardiomyocyte Polyploidy and Implications for Heart Regeneration. Annu. Rev. Physiol. 82, 45–61 (2020).

7. Mollova, M. et al. Cardiomyocyte proliferation contributes to heart growth in young humans. Proc. Natl. Acad. Sci. U. S. A. 110, 1446–1451 (2013).

8. Donne, R., Saroul-Aïnama, M., Cordier, P., Celton-Morizur, S. & Desdouets, C. Polyploidy in liver development, homeostasis and disease. Nat. Rev. Gastroenterol. Hepatol. 17, 391–405 (2020).

9. Ravid, K., Lu, J., Zimmet, J. M. & Jones, M. R. Roads to polyploidy: The megakaryocyte example. J. Cell. Physiol. 190, 7–20 (2002).

10. Yagi, M. et al. DC-STAMP is essential for cell-cell fusion in osteoclasts and foreign body giant cells. J. Exp. Med. 202, 345–351 (2005).

11. Sladky, V. C., Eichin, F., Reiberger, T. & Villunger, A. Polyploidy control in hepatic health and disease. J. Hepatol. 75, 1177–1191 (2021).

12. Kiermaier, E., Stötzel, I., Schapfl, M. A. & Villunger, A. Amplified centrosomes—more than just a threat. EMBO Rep. 25, 4153–4167 (2024).

13. Tinel, A. & Tschopp, J. The PIDDosome, a Protein Complex Implicated in Activation of Caspase-2 in Response to Genotoxic Stress. Science (80-. ). 304, 843–846 (2004).

14. Tinel, A. et al. Autoproteolysis of PIDD marks the bifurcation between pro-death caspase-2 and pro-survival NF-κB pathway. EMBO J. 26, 197–208 (2007).

15. Janssens, S., Tinel, A., Lippens, S. & Tschopp, J. PIDD Mediates NF-κB activation in response to DNA damage. Cell 123, 1079–1092 (2005).

16. Ando, K. et al. PIDD Death-Domain Phosphorylation by ATM Controls Prodeath versus Prosurvival PIDDosome Signaling. Mol. Cell 47, 681–693 (2012).

17. Burigotto, M. et al. Centriolar distal appendages activate the centrosome-PIDDosome-p53 signalling axis via ANKRD26. EMBO J. 1–22 (2020) doi:10.15252/embj.2020104844.

18. Fava, L. L. et al. The PIDDosome activates p53 in response to supernumerary centrosomes. Genes Dev. 31, 34–45 (2017).

19. Evans, L. T. et al. ANKRD26 recruits PIDD1 to centriolar distal appendages to activate the PIDDosome following centrosome amplification. EMBO J. 1–18 (2020) doi:10.15252/embj.2020105106.

20. Garcia-Carpio, I. et al. Extra centrosomes induce PIDD1 -mediated inflammation and immunosurveillance. EMBO J. 43, 1–22 (2023).

21. Bouchier-Hayes, L. & Green, D. R. Caspase-2: The orphan caspase. Cell Death Differ. 19, 51–57 (2012).

22. Oliver, T. G. et al. 2011 【Mol. Cell.】 Caspase-2-mediated cleavage of Mdm2 creates p53-induced.pdf. 43, 57–71 (2012).

23. Kim, J. Y. et al. ER Stress Drives Lipogenesis and Steatohepatitis via Caspase-2 Activation of S1P. Cell 175, 133–145.e15 (2018).

24. Billen, L. P., Shamas-Din, A. & Andrews, D. W. Bid: a Bax-like BH3 protein. Oncogene 27, S93–S104 (2008).

25. Rizzotto, D. et al. Caspase-2 kills cells with extra centrosomes. Sci. Adv. 10, eado6607 (2024).

26. Sladky, V. C. et al. E2F-Family Members Engage the PIDDosome to Limit Hepatocyte Ploidy in Liver Development and Regeneration. Dev. Cell 52, 335–349.e7 (2020).

27. Shah, R. B. et al. FANCI functions as a repair/apoptosis switch in response to DNA crosslinks. Dev. Cell 56, 2207–2222.e7 (2021).

28. Shah, R. B., Li, Y., Yu, H., Kini, E. & Sidi, S. Stepwise phosphorylation and SUMOylation of PIDD1 drive PIDDosome assembly in response to DNA repair failure. Nat. Commun. 15, 9195 (2024).

29. Ando, K. et al. NPM1 directs PIDDosome-dependent caspase-2 activation in the nucleolus. J. Cell Biol. 216, 1795–1810 (2017).

30. Brown-Suedel, A. N. & Bouchier-Hayes, L. Caspase-2 Substrates: To Apoptosis, Cell Cycle Control, and Beyond. Front. Cell Dev. Biol. 8, 1–17 (2020).

31. Logette, E. et al. PIDD orchestrates translesion DNA synthesis in response to UV irradiation. Cell Death Differ. 2011 186 18, 1036–1045 (2011).

32. Tinel, A. et al. Regulation of PIDD auto-proteolysis and activity by the molecular chaperone Hsp90. Cell Death Differ. 18, 506–515 (2011).

33. Tinel, A. The PIDDosome, a Protein Complex Implicated in Activation of Caspase-2 in Response to Genotoxic Stress. 304, 843–847 (2004).

34. Burigotto, M. et al. Centriolar distal appendages activate the centrosome-PIDDosome-p53 signalling axis via ANKRD26. EMBO J. 40, 1–22 (2021).

35. Evans, L. T. et al. ANKRD26 recruits PIDD1 to centriolar distal appendages to activate the PIDDosome following centrosome amplification. EMBO J. 40, 1–18 (2021).

36. Kim, J. Y. et al. PIDDosome-SCAP crosstalk controls high-fructose-diet-dependent transition from simple steatosis to steatohepatitis. Cell Metab. 34, 1548–1560.e6 (2022).

37. Hörst, F. et al. CellViT: Vision Transformers for precise cell segmentation and classification. Med. Image Anal. 94, 103143 (2024).

38. Leone M, Kinz N, Eichin F, Obwegs D, Sladky VC, Braun VZ, Rizzotto D, E. L. & Manzl C, Moos K, Julia Mergner, Giansanti P, Garcia MN, Marques MM, Jacotot ED, Savko C, Boerries M, Sussman MA, V. A. The PIDDosome controls cardiomyocyte polyploidization during 1 postnatal heart development. bioRxiv (2025) doi:10.1101/2024.08.27.609375.

39. Zebrowski, D. C. et al. Developmental alterations in centrosome integrity contribute to the post-mitotic state of mammalian cardiomyocytes. Elife 4, (2015).

40. Redecke, V. et al. Hematopoietic progenitor cell lines with myeloid and lymphoid potential. Nat. Methods 10, 795–803 (2013).

41. Braun, V. Z. et al. Extra centrosomes delay DNA damage–driven tumorigenesis. Sci. Adv. 10, 1–16 (2024).

42. Montes, R., Luna, D. O., Wagner, D. S. & Lozanot, G. Rescue of early embryonic lethality in mdm2·deficient mice by deletion of. 378, 203–206 (1995).

43. Jones, S. N., Roe, A. E., Donehowert, L. A. & Bradley, A. Rescue of embryonic lethality in Mdm2-deficient mice by absence of p53. 378, 847–849 (1995).

44. Kim, J. Y. et al. PIDDosome-SCAP crosstalk controls high-fructose-diet-dependent transition from simple steatosis to steatohepatitis. Cell Metab. 34, 1548–1560.e6 (2022).

45. Ran, F. A. et al. Genome engineering using the CRISPR-Cas9 system. Nat. Protoc. 8, 2281–2308 (2013).

46. Grompe, M., Jones, S. N., Loulseged, H. & Caskey, C. T. Retroviral-Mediated Gene Transfer of Human Ornithine Transcarbamylase into Primary Hepatocytes of spf and spf-ash Mice. Hum. Gene Ther. 3, 35–44 (1992).

47. LazySlide: Accessible and interoperable whole slide image analysis — LazySlide 0.6.0 documentation. https://lazyslide.readthedocs.io/en/latest/.

48. Chen, R. J., et al. Towards a general-purpose foundation model for computational pathology. Nature Medicine vol. 30 (Springer US, 2024).

49. Charlier, F. et al. trevismd/statannotations: v0.5. (2022) doi:10.5281/zenodo.7213391.

