## Supplementary figures and images for "Sequential PIDD1 auto-processing is essential for ploidy control in the liver and heart"

### suppl Fig

Suppl. Figure 1

A

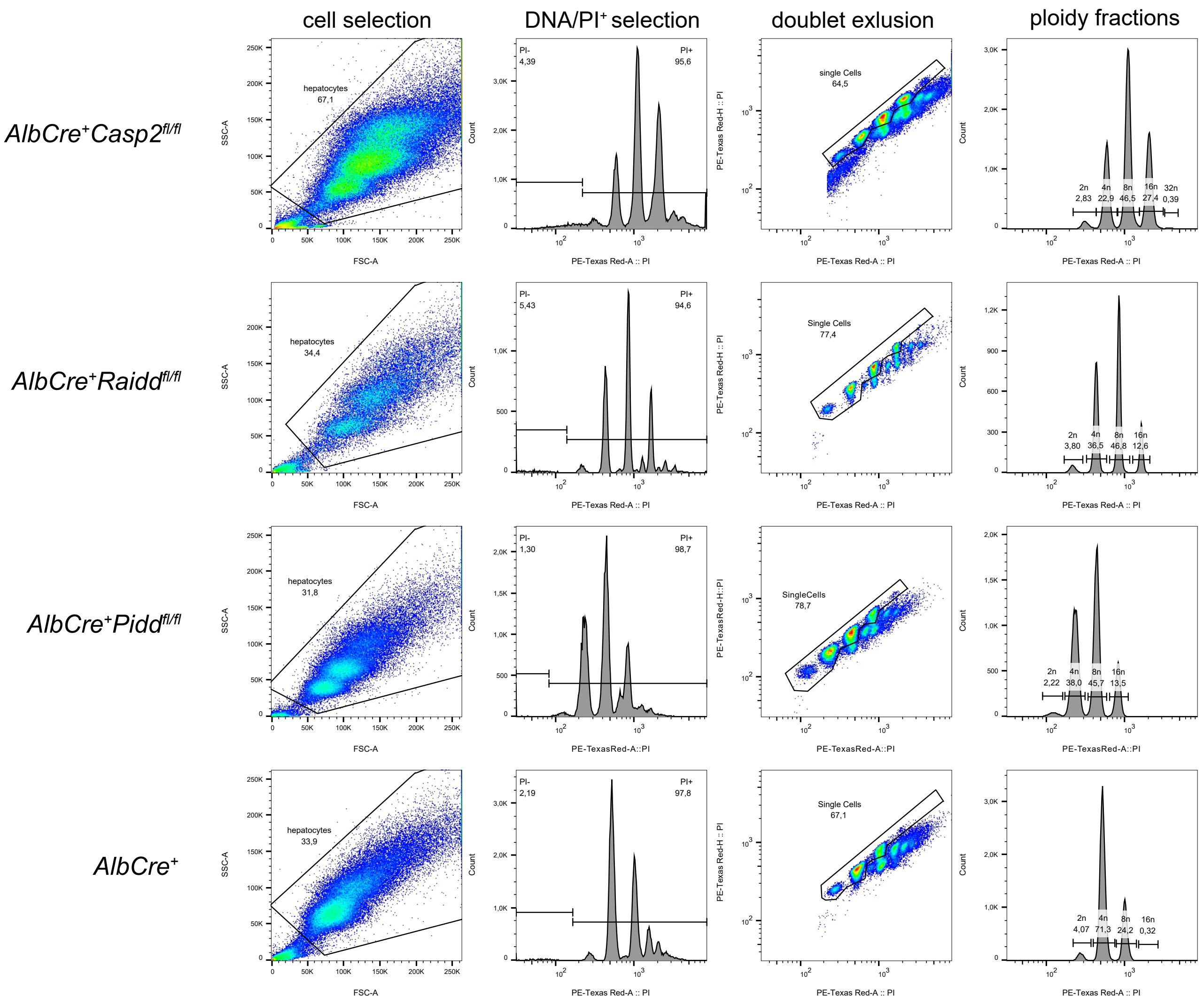

Suppl. Figure 2

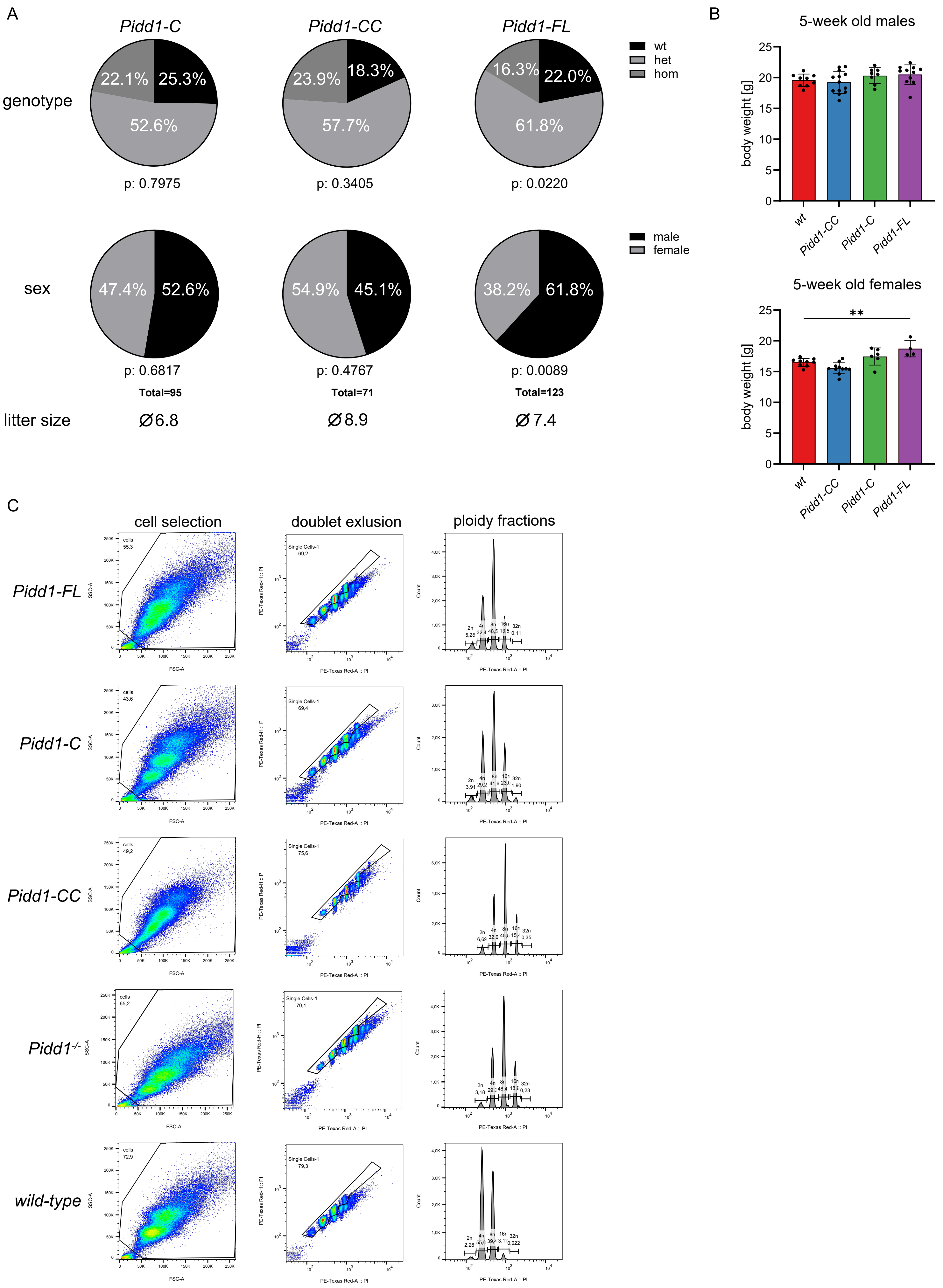

Suppl. Figure 3

A

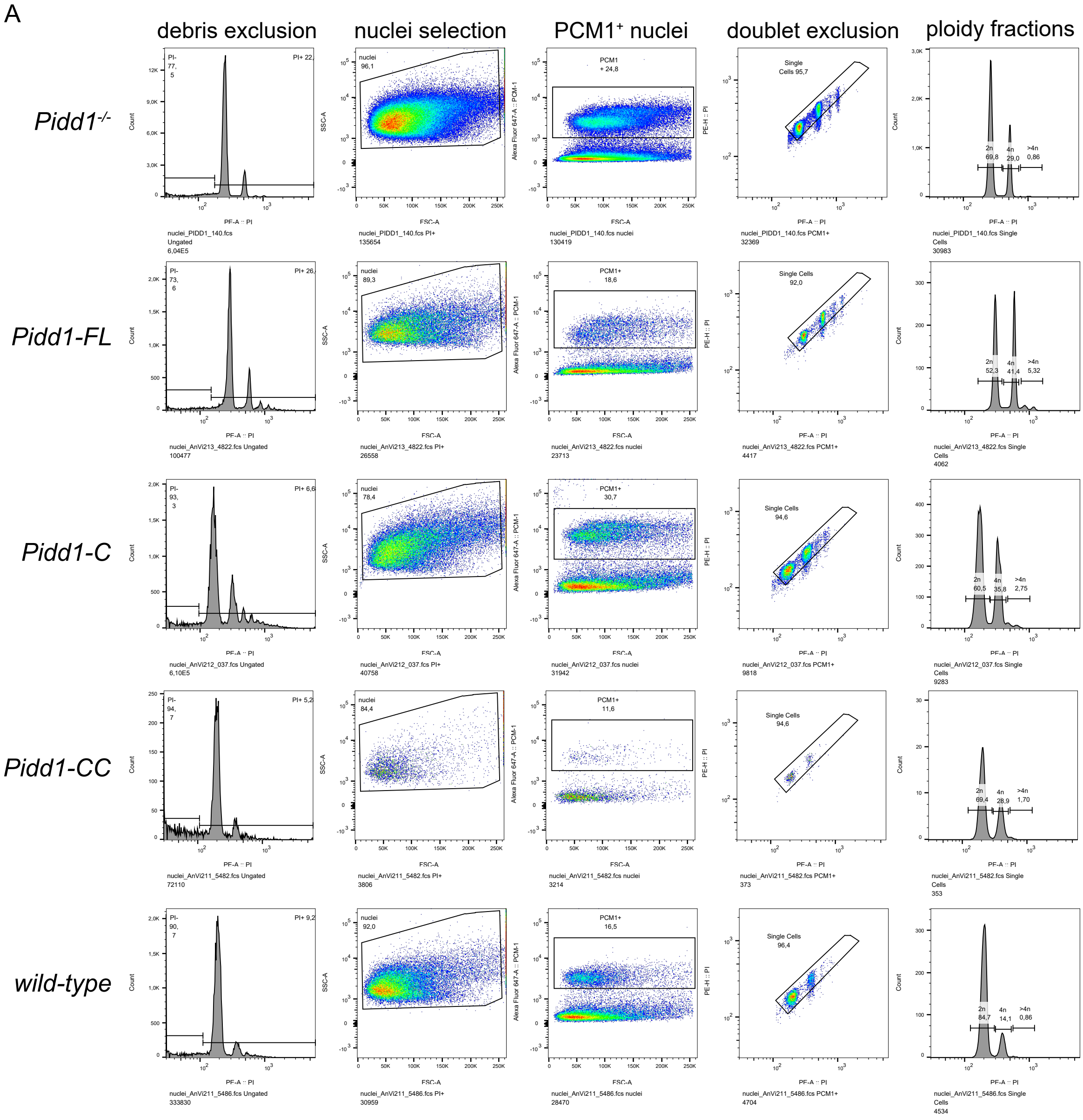

Suppl. Figure 4

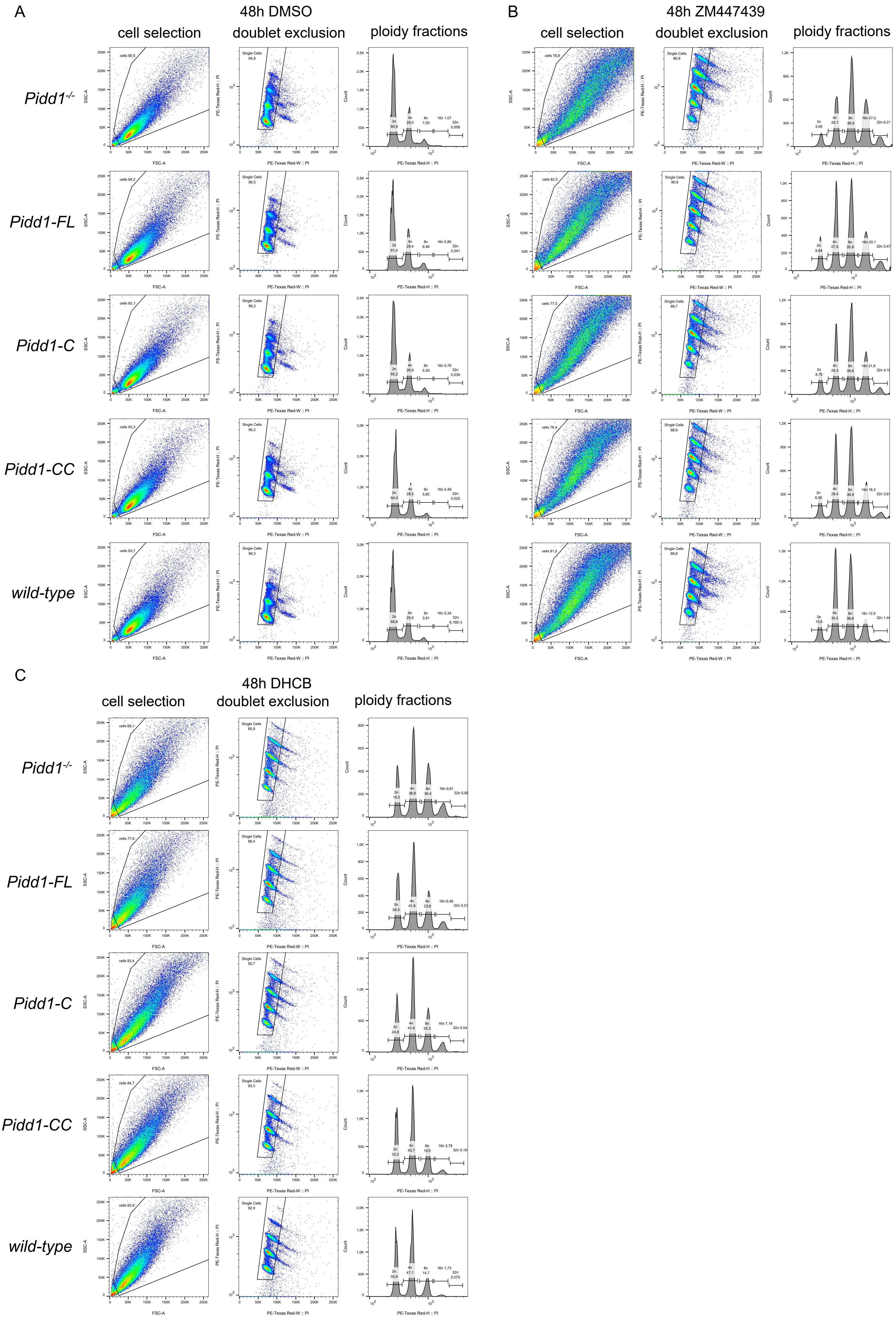

Suppl. Figure 5

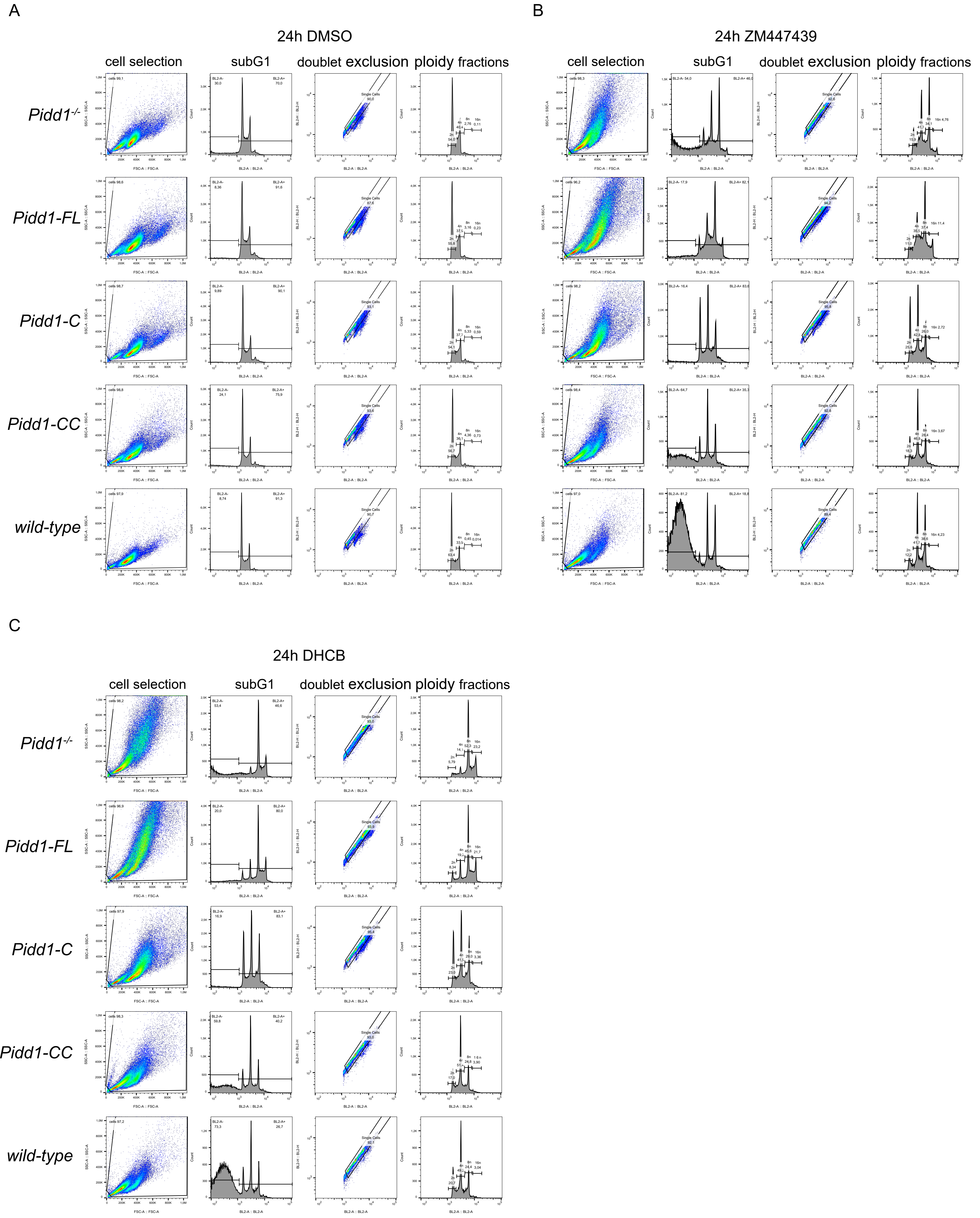

Suppl. Figure 6

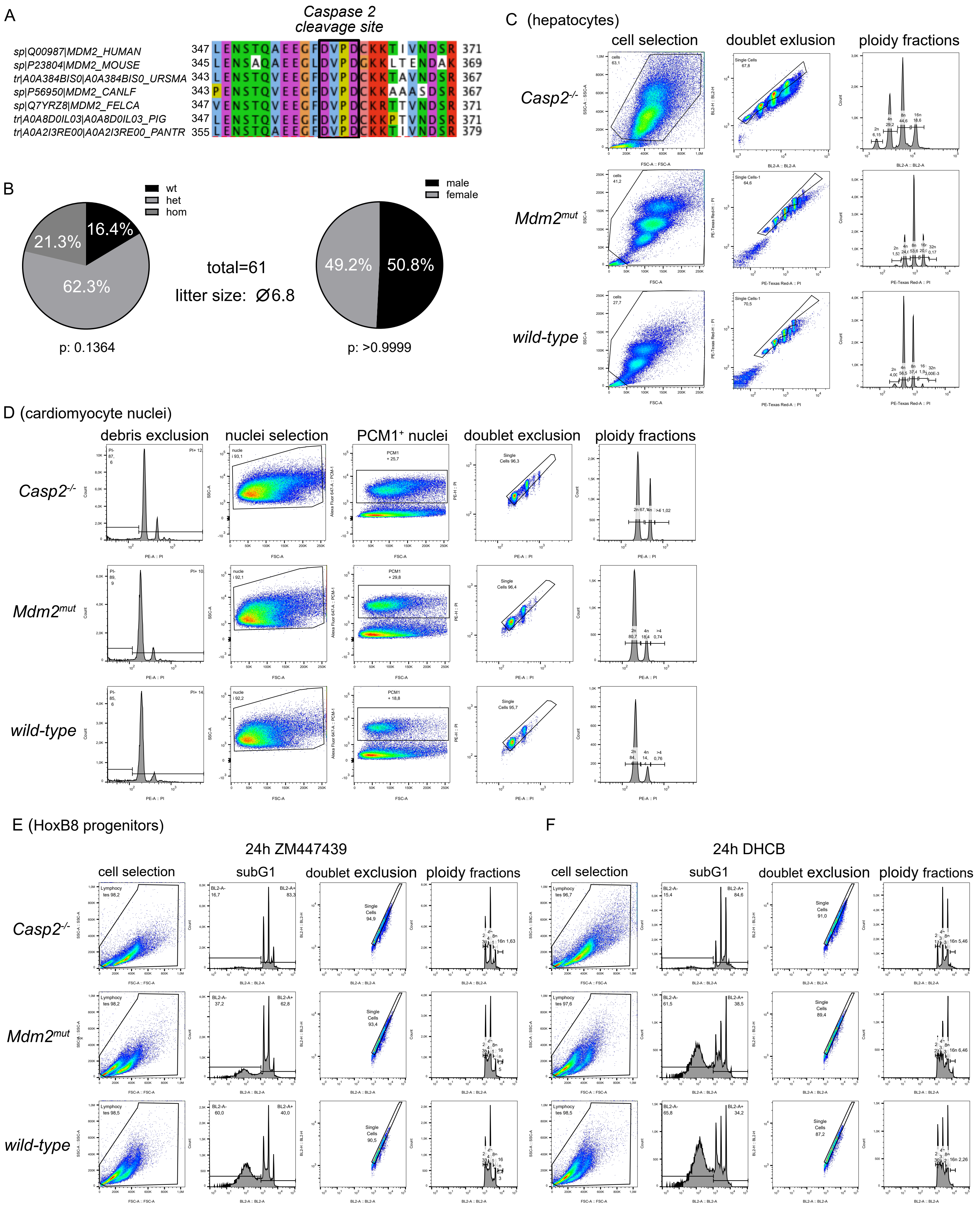
